# Reconstructing signaling history of single cells with imaging-based molecular recording

**DOI:** 10.1101/2024.10.11.617908

**Authors:** Kai Hao, Mykel Barrett, Zainalabedin Samadi, Amirhossein Zarezadeh, Yuka McGrath, Amjad Askary

## Abstract

The intensity and duration of biological signals encode information that allows a few pathways to regulate a wide array of cellular behaviors. Despite the central importance of signaling in biomedical research, our ability to quantify it in individual cells over time remains limited. Here, we introduce INSCRIBE, an approach for reconstructing signaling history in single cells using endpoint fluorescence images. By regulating a CRISPR base editor, INSCRIBE generates mutations in genomic target sequences, at a rate proportional to signaling activity. The number of edits is then recovered through a novel ratiometric readout strategy, from images of two fluorescence channels. We engineered human cell lines for recording WNT and BMP pathway activity, and demonstrated that INSCRIBE faithfully recovers both the intensity and duration of signaling. Further, we used INSCRIBE to study the variability of cellular response to WNT and BMP stimulation, and test whether the magnitude of response is a stable, heritable trait. We found a persistent memory in the BMP pathway. Progeny of cells with higher BMP response levels are likely to respond more strongly to a second BMP stimulation, up to 3 weeks later. Together, our results establish a scalable platform for genetic recording and *in situ* readout of signaling history in single cells, advancing quantitative analysis of cell-cell communication during development and disease.

## Introduction

In multicellular organisms, the behavior and fate of cells are orchestrated by signaling pathways. Cells interpret the intensity and duration of these signals to produce the appropriate response. For example, during development, signaling gradients provide spatial and temporal cues that guide cell fate decisions ^1–3^. Immune response is also coordinated by cell-cell signaling; the amplitude and duration of signals from the T cell receptor can determine whether a T cell activates, proliferates, and differentiates, or becomes tolerant ^4^. Similarly, neoplastic cell division and cancer metastasis often involve aberrant signaling levels. WNT signaling is frequently up-or down-regulated in different types of cancer, and the pathway activity level can serve as a prognostic indicator of patient outcomes ^5^. In all these contexts, explaining or predicting cellular behavior hinges on accurately measuring the activity of signaling pathways in individual cells over time.

The significance of cell signaling has spurred development of several strategies for capturing the molecular history of cells ^1,6^. Time lapse microscopy is the most direct approach. It typically involves reporters that express a fluorescent protein at a level proportionate to the amount of signaling activity. Time lapse imaging provides superb temporal resolution, however, it is limited to optically accessible samples, and relatively short time scales. There is also a trade off between temporal resolution and the size of the sample that can be effectively analyzed ^7^. Despite significant progress in microscopy techniques, it remains difficult to reliably track large numbers of cells in complex *in vivo* environments over extended time periods.

Genetic engineering provides an alternative to direct observation. Using site-specific recombination, cells can be permanently labeled upon activation of a signaling pathway. This powerful technique has enabled numerous groundbreaking discoveries ^8^. However, it seldom offers the precision and spatiotemporal resolution required for quantitative association of signaling levels and cellular phenotypes. Site-specific recombination typically has a binary outcome: cells that activated the recombinase at high enough level to catalyze the recombination versus those that did not.

Molecular recording offers a promising solution to these challenges by engineering cells to record their signaling history in their genome, as targeted mutations that can be detected at a later time (Figure 1A). This approach is enabled by the multitude of genome editing methods, including recombinases^9–11^, CRISPR/Cas9^12–16^, CRISPR intergrases ^17–19^, base editors^20–22^, and prime editors^23–25^, which can alter the sequence of genomic DNA in living cells. The pattern and frequency of mutations can be used to reconstruct molecular history of the cells, as well as lineage relationship between them ^26–29^. Recording longitudinal information in the genome allows researchers to use high-throughput snapshot methods, such as single cell sequencing and spatial genomics, to capture history of the cells as well as their endpoint state. It also circumvents the need for direct real time observation of the system.

**Figure 1.**
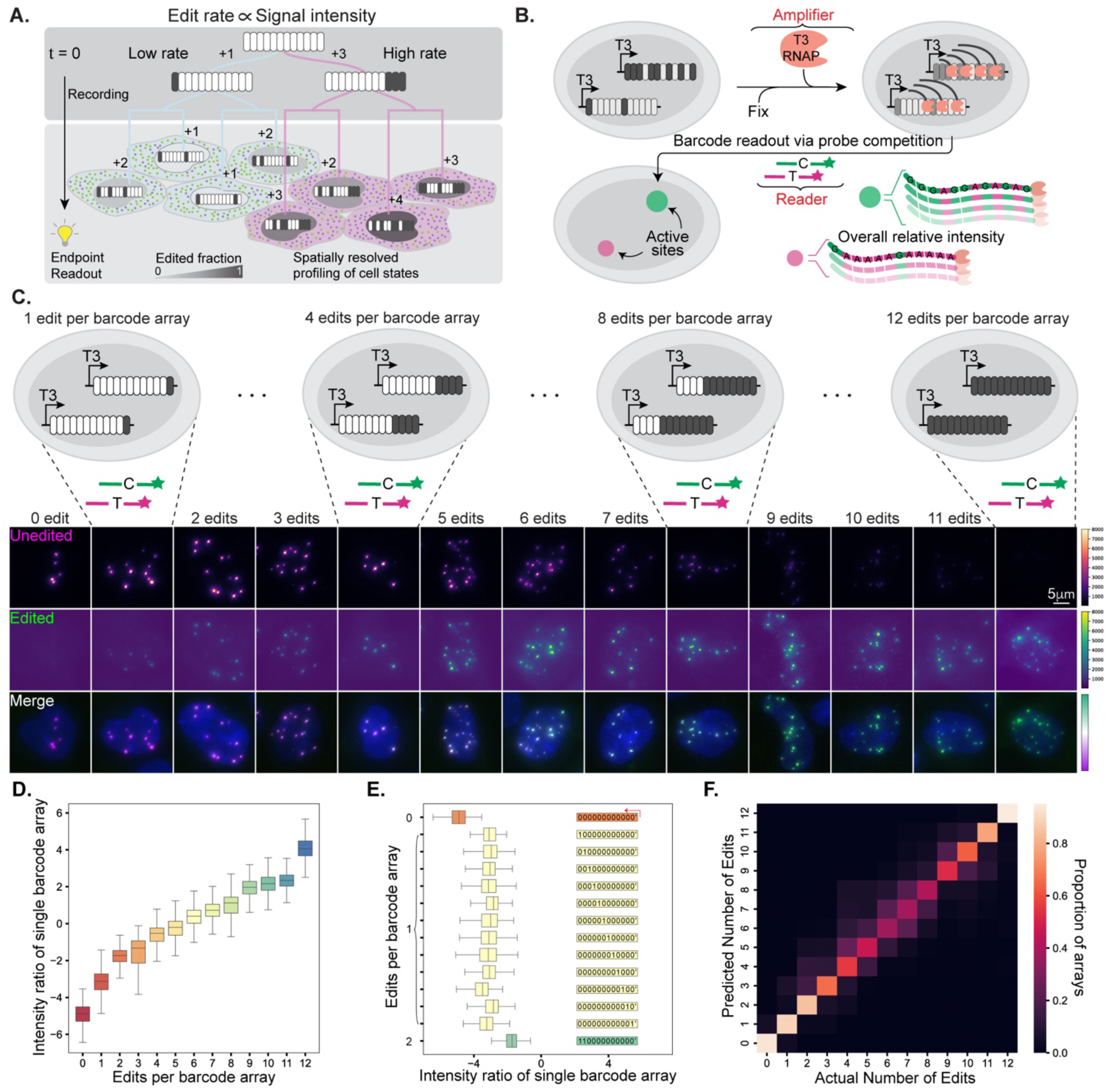
Molecular recording with in situ ratiometric barcode readout. **(A)** Each barcode is a genomic site that can be edited to provide two possible states, represented by white and gray ovals. When edit rate is proportional to signal intensity, the number of edited barcodes maintain a record of signaling activity in the lineage of each cell. Barcode edits can be read out together with spatial information and gene expression profile of the cells at the endpoint. The nuclei are shaded in varying grayscales, representing the fraction of edited barcodes in each nucleus. **(B)** Workflow of ratiometric barcode readout. Barcode arrays are transcribed in situ, using recombinant T3 RNA polymerase. Accumulation of transcripts in the active site amplifies the signal. Barcode transcripts are then probed with an equimolar mixture of two fluorescently labeled probes, corresponding to the edited and unedited states (C and T respectively). **(C)** Representative images of cells with known edit states. Each column shows one edit level, with the fluorescence channel corresponding to unedited and edited probes, in magenta and green, respectively. As the number of edited barcodes in the arrays increases from 0 to 12 (left to right), signal in the edited channel increases and signal in the unedited channel diminishes. **(D)** Intensity ratio, defined as log2-scaled ratio between the fluorescence intensity in the edited and unedited channels, increases with increasing number of edits in the barcode arrays. **(E)** Regardless of the position of the edited barcode within the array, arrays with one edit show similar intensity ratios, which are higher than the array with no edit and lower than the array with two edits. ‘1’ stands for edited barcode, ‘0’ stands for unedited barcode, and the red arrow indicates the direction of T3 transcription. **(F)** The number of edits in barcode arrays can be inferred from fluorescence images of two competing probes. Confusion matrix compares the true edit numbers against the predicted outputs from the CNN model. The frequency values are normalized so that each row sums to one. All results in this figure are from analysis of the BC1 array.

An ideal system for genetically recording signaling activity should provide accurate single-cell measurements, be compatible with collecting additional information (such as gene expression and spatial context), have the potential to simultaneously record multiple signals, and be user-friendly without requiring specialized tools or equipment. Existing methods that use next generation sequencing for readout, while promising, disrupt spatial organization, which is often essential for understanding tissue biology and function. Further, due to the digital and probabilistic nature of DNA editing, a large number of memory elements are required to achieve robust analog recording. This is accomplished either through integration of multiple barcode arrays in each cell ^29^, or by targeting several endogenous sites ^11,19,21^.

However, it is difficult to consistently recover both the transcriptional profile and the sequence of multiple genomic targets from individual cells by single cell sequencing, as this occurs infrequently ^23,30,31^. Consequently, existing molecular recording strategies have limited accuracy at the single-cell level.

Imaging-based methods preserve spatial information, can decode numerous target sites in each cell ^22^, and are compatible with gene expression profiling ^10^. However, performing the many rounds of hybridization and imaging that is required to read out the recorded information is challenging. Sequential imaging is also time consuming, limiting the total sample area that can be analyzed in a reasonable timeframe. As a result, application of imaging-based molecular recording has remained limited.

Here, we introduce a novel approach for detecting the number of edits in DNA barcodes based on the fluorescence intensity of only two probes. This approach, which we call ratiometric barcode readout, bypasses the sequential hybridization and imaging process, thereby solving the main hurdle for widespread application of imaging-based molecular recording. Using ratiometric barcode readout, we developed a scalable method for spatially resolved quantitative reconstruction of signaling activity at the single cell level, called, <u>IN</u> situ <u>S</u>ingle <u>C</u>ell <u>R</u>ecording of signal Intensity as <u>B</u>arcode <u>E</u>dits (INSCRIBE). INSCRIBE uses a CRISPR A-to-G base editor (ABE) to mutate specific sites within synthetic barcode arrays that are distributed across the genome. Using signal responsive cis-regulatory elements, we made Human Embryonic Kidney (HEK293) cell lines in which frequency of barcode edits is proportional to activity of WNT or BMP signaling pathways. We show that the amplitude and duration of signaling in these cell lines can be recovered from endpoint measurements that involve only a single round of imaging.

Using INSCRIBE, we analyzed single cell variability in response to WNT and BMP stimulation. Even in a clonal population, individual cells can exhibit varying levels of response, although they are exposed to the same level of inducer. We asked whether this heterogeneity reflects stable and heritable differences between cells. Comparing the magnitude of response between two sequential stimulations revealed a persistent memory in the BMP pathway, which can last up to 18 days. This study demonstrates the utility of INSCRIBE in providing novel biological insight and addressing otherwise intractable questions.

## Results

### Ratiometric readout efficiently recovers barcode array states along with spatial information

Molecular recording involves engineering cells to accumulate mutations in engineered genomic targets, referred to here as “barcode arrays”. If the mutation rate is proportional to the activity of a signaling pathway, the fraction of edited barcodes reflects the cumulative pathway activity in the lineage of each cell **(Fig. 1A)**. As the foundation of a scalable and broadly accessible imaging-based molecular recording system, we sought to develop a simple method to recover the fraction of edited barcodes in barcode arrays.

We recently developed a method for sensitive *in situ* readout of DNA barcodes ^32^. This strategy uses a phage RNA polymerase to transcribe the barcode arrays in situ and capture the resulting transcripts in the active site of transcription, which can be visualized as bright spots by Fluorescence *In Situ* Hybridization (FISH). Hybridization with competing probes can then enable accurate *in situ* identification of single nucleotide edits. For barcodes with two possible states, edited and unedited, this approach would require two fluorescence channels for each barcode. Since dozens of barcodes are required to accurately infer the signaling history of a cell, decoding the state of each barcode separately would require numerous rounds of hybridization and imaging. We argued that the information needed to infer signaling activity level can be obtained much more efficiently if all barcodes in an array are probed with the same pair of probes against the edited and unedited states **(Fig. 1B)**. This approach, which we call ratiometric barcode readout, detects the fraction of edited barcodes in an array, instead of the state of each individual barcode. So, it trades off acquiring unnecessary information for simplicity and scalability.

We assessed the feasibility of ratiometric barcode readout using cell lines that contain synthetic barcode arrays with known states. Our barcode arrays are designed to include 12 target sites for a single guide RNA (gRNA), downstream of T3 promoter **(Fig. 2A)**. Each target site has only one A nucleotide within the editing window of A-to-G base editor (ABE) ^33,34^, which overlaps with the probe sequence we use for FISH **(Fig. 2B)**. We designed two independent barcode arrays which are targeted by orthogonal gRNAs, henceforth referred to as BC1 **(Fig. 1C-F)** and BC2 **(Fig. S1)**. We created DNA constructs containing barcode arrays with 0 to 12 edits, in different configurations, flanked with piggyBac inverted terminal repeats (ITRs) to enable genomic integration. We then made HEK293 cell lines, each containing multiple stable integrations for one of these synthetic barcode array constructs.

**Figure 2.**
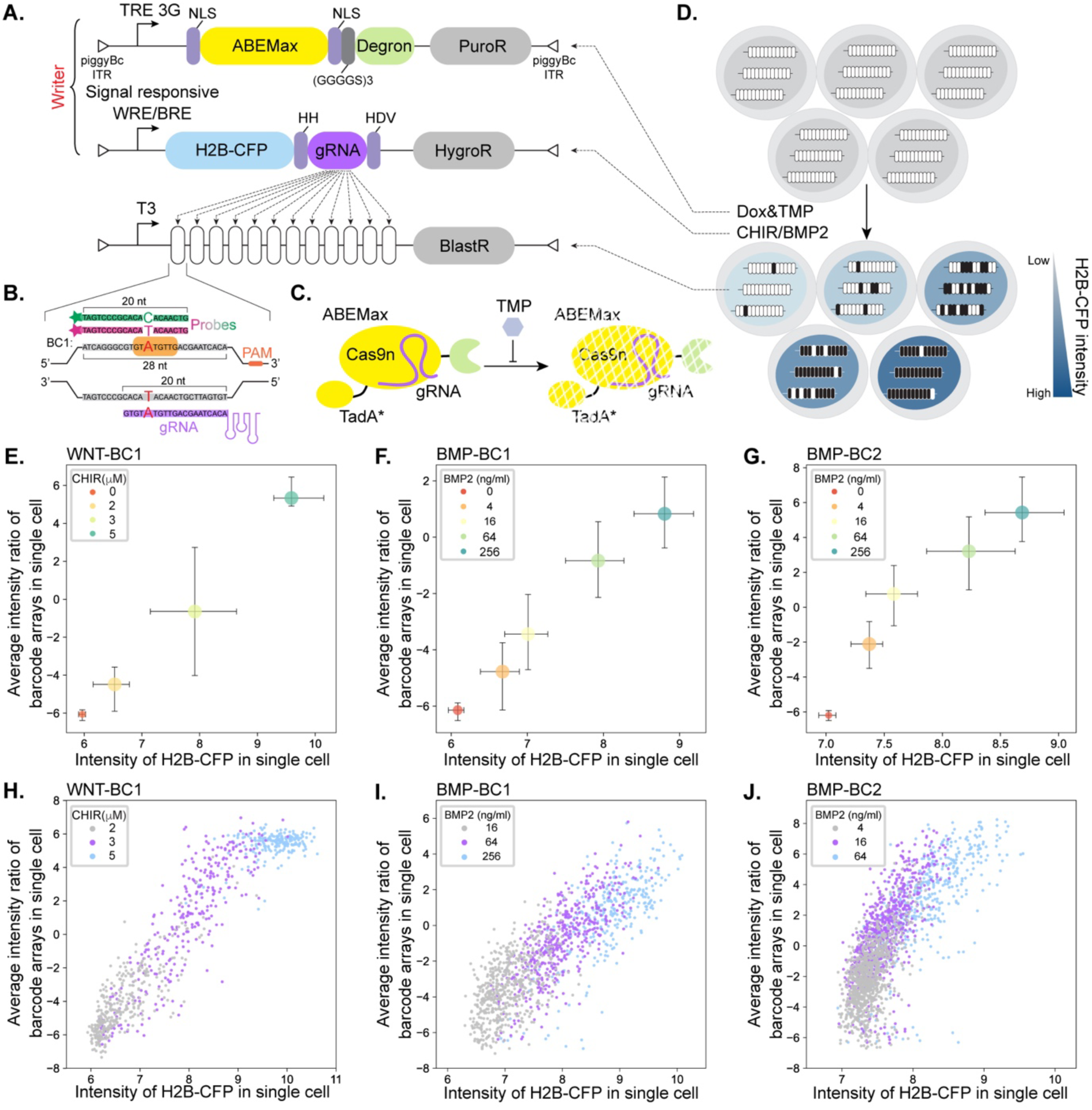
Recording the activity of WNT and BMP pathways in a dose dependent manner. **(A)** Design of the INSCRIBE cell lines. Each cell contains doxycycline (Dox) inducible ABE, signal responsive gRNA and H2B-CFP, and barcode arrays compatible with ratiometric readout. All three components are stably integrated using piggyBac transposase. **(B)** Each barcode consists of a gRNA target site that partially overlaps with the probe target sequence. The sequence of a single barcode in the BC1 array with its corresponding gRNA target site and the probe binding site are shown. The editing window of ABE is shaded in orange. Each barcode array contains 12 identical barcodes. Sequences of the BC2 array are listed in Supplementary table 1. **(C)** ABE expression is further controlled through stabilization of ecDHFR degron by trimethoprim (TMP). **(D)** The WNT and BMP pathways are activated by CHIR and BMP2, respectively, in the corresponding INSCRIBE cell lines. Signal dependent expression of gRNA leads to accumulation of edits in barcode arrays. It is also accompanied by expression of H2B-CFP, which is used as a proxy for signaling activity in short term (3 day) recording experiments. **(E, F and G)** Scatter plots display the population median of single-cell barcode edit levels versus H2B-CFP intensity for WNT-BC1 **(E)**, BMP-BC1 **(F)**, and BMP-BC2 **(G)** cells. Edit level in each cell is calculated as the average of intensity ratios for all its barcode arrays. The recording was conducted over three days with varying levels of signal inducer, indicated by color coding. Bars represent the interquartile ranges. **(H, I and J)** Barcode edit levels in individual cells are correlated with their H2B-CFP intensities. Each point in the scatter plots represents a single cell, with colors indicating the level of the inducer. ABE expression was induced by 2500 ng/ml Dox for WNT recording **(E and H)** and 500 ng/ml Dox for BMP recording **(F, I, G, and J)**.

After *in situ* transcription by T3 RNA polymerase, using our previously described approach ^32^, we hybridized two fluorescently labeled probes, complementary to the edited and unedited states of the target sites. The resulting images revealed a trend: as the number of edits increased, fluorescence intensity in the channel for the edited state rose, while intensity in the channel for the unedited state diminished **(Fig. 1C and Fig. S1A)**. This effect can be shown quantitatively, by calculating the ratio of fluorescence intensity in the edited versus unedited channel **(Fig. 1D and Fig. S1B)**. Comparing barcode arrays with the same number of edits, but in different configurations, showed that intensity ratio is a function of the number of edits, not their position within the array **(Fig. 1E and Fig. S1C)**.

Knowing the true state of each barcode array in these experiments allows us to train a classifier to predict the number of edits from the microscopy image of an array. We trained a Convolutional Neural Network (CNN) for this task, using images of 23,743 and 15,855 spots, for BC1 and BC2 respectively, corresponding to barcode arrays with 0 to 12 edits **(Fig. 1F and Fig. S1D)**. The trained classifier was able to infer the number of edits in barcode arrays with a macro-averaged F1 score of 0.6 for BC1 and 0.69 for BC2. Accuracy was greater for arrays with a low (0 to 3) or high (10 to 12) number of edits, while the majority of inaccurate predictions occurred in arrays with intermediate edit counts. In total, 91.5% of BC1 arrays and 97.7% of BC2 arrays had predicted edit counts within 2 edits of their actual counts. This confirms that ratiometric barcode readout can infer the edit level of barcode arrays from two-channel images and provides an automated classification tool for this purpose.

### INSCRIBE enables analog recording of signaling amplitude in single cells

To demonstrate genetic recording of signaling activity with single cell imaging-based readout, we established monoclonal HEK293 cell lines harboring three components: an inducible ABE, a signal responsive gRNA, and barcode arrays compatible with ratiometric readout **(Fig. 2A-B)**. ABE expression in INSCRIBE cell lines can be controlled by doxycycline (Dox) using the Tet-On 3G system ^35^. To minimize the possibility of basal editing, ABE is also fused to an ecDHFR degron sequence which can be stabilized by trimethoprim (TMP) ^36^ **(Fig. 2C)**. Edit rate is coupled to either BMP or WNT pathway activity using cis-regulatory elements (CREs) containing multimerized binding sites for BMP SMADs^37^ and TCF/LEF ^38,39^, respectively. These CREs drive signal dependent production of a transcript encoding both Cyan Fluorescent Protein fused to Histone H2B (H2B-CFP) and a barcode specific gRNA **(Fig. 2A)**. gRNA is released from the transcript using Hammerhead (HH) and Hepatitis Delta Virus (HDV) ribozymes ^40,41^. To test the performance of different barcode arrays, we made BMP recorder cell lines with either BC1 or BC2. WNT recorder lines were made only with BC1.

Constructs were stably integrated in HEK293 cells with three rounds of piggyBac transposition **(Methods)**. We then recovered and analyzed multiple monoclonal cell lines with all INSCRIBE components. BMP signaling was induced by adding 64 ng/ml of recombinant BMP2 and the WNT pathway was stimulated by 3 µM of CHIR99021 (CHIR), a small molecule GSK-3 inhibitor. In addition, the culture media contained 100 ng/ml of Dox and 10 µM TMP. Different clones showed some variability in their baseline activity and sensitivity to stimulation **(Fig. S2)**. For all subsequent experiments, we focused on one clone for each pathway-barcode array pair that had minimal baseline activity and relatively homogenous response after stimulation.

To assess whether INSCRIBE could record signaling activity of BMP and WNT pathways, we exposed the engineered cells to varying amounts of BMP2 and CHIR, respectively. After culturing the cells for 3 days, we performed *in situ* T3 transcription and hybridized the samples with two fluorescently labeled probes for the edited and unedited states. We then imaged cells in the two channels associated with the probes, as well as DAPI to label nuclei, and CFP. The reporter for each pathway drives the expression of a chimeric transcript that cleaves into H2B-CFP mRNA and barcode targeting gRNA. Since H2B-CFP protein is relatively stable, it is mainly diluted by cell division that takes approximately 24 to 48 hours. Thus, for short recording experiments (3 days in this case) accumulation of H2B-CFP provides a good proxy for the reporter activity in each cell **(Fig. 2D)**.

In the resulting images, we quantified the editing level of each cell as the average of intensity ratio for all its barcode arrays **(Methods)**. For both the BMP and WNT pathways, the edit level of cell populations increases with increasing concentrations of pathway inducers, in a manner correlated with CFP level **(Fig. 2E-G)**. More importantly, this trend was also observed at the single cell level **(Fig. 2H-J)**, demonstrating the ability of INSCRIBE to recover the signaling history of individual cells. Further, BC1 and BC2 showed different dynamics in response to BMP2. BMP-BC1 cells had lower edit levels compared to BMP-BC2 cells cultured in the same concentration of BMP2 **(Fig. S2D)**. As a result, higher concentrations of BMP2 (16, 64, and 256ng/ml) led to distinct edit levels in BMP-BC1 cells **(Fig. 2I)**. Whereas, BMP-BC2 cells had distinguishable edit levels for lower concentrations of BMP2 (4, 16, and 64ng/ml), with 256ng/ml of BMP2 leading to saturation **(Fig. 2G and 2J)**. This effect is likely due to differences in the efficiency of gRNAs targeting each barcode array. Therefore, we expect different barcode arrays to have different dynamic ranges, suitable for recording signals with different amplitudes.

We also asked whether the dynamic range of INSCRIBE can be tuned by regulating ABE. Varying Dox levels from 100 to 2500 ng/ml did not have a significant effect on edit levels in any of the tested conditions **(Fig. S3)**. At 20 ng/ml, ABE was not induced sufficiently for recording BMP pathway activity, while there were detectable edit levels in response to 5 µM CHIR. Therefore, regulation of ABE expression appears to have a binary outcome, with edit levels being a function of signaling activity once ABE is present at high enough levels.

Together, our results here show that INSCRIBE is an analog recording system capable of recovering signaling amplitude over a fixed time window from single cells. The sensitivity of INSCRIBE can be adjusted using different gRNAs. Recording works over a wide range of ABE levels, facilitating implementation of the system in other cell lines or *in vivo* models.

### INSCRIBE can record duration of signaling activity

In addition to intensity, duration of signaling activity is crucial in regulating cellular behavior ^42^. Fluorescent reporters dilute over time by degradation and cell division. Therefore, it is difficult to estimate the signaling duration from their endpoint level. In contrast, barcode edits are irreversible and inheritable, making them ideal for recording signaling history. To test if INSCRIBE can recover signaling duration, we cultured cells for different amounts of time under editing conditions (500 ng/ml Dox and 10 µM TMP) combined with 64ng/ml BMP2 or 3µM CHIR **(Fig. 3A)**. We used our CNN classifier to analyze endpoint images and estimate the average edit level in each cell after determining the number of edits in each barcode array (Methods). As anticipated, barcode edits accumulated progressively over time in all cases **(Fig. 3B)**. In BMP-BC1 and WNT-BC1 cells, this increase was nearly monotonic over a span of 7 days. In contrast, BMP-BC2 cells exhibited a rapid increase in edits within the first 3 days, nearing a fully edited state after 4 days. This is consistent with the faster edit rate of BC2 array **(Fig. S2D)**. Together, our results demonstrate that INSCRIBE can record either the amplitude of a signal over a defined time period **(Fig. 2)** or the duration of a signal of known intensity **(Fig. 3)**, all in a format that is image-readable and can be recovered from single cells.

**Figure 3.**
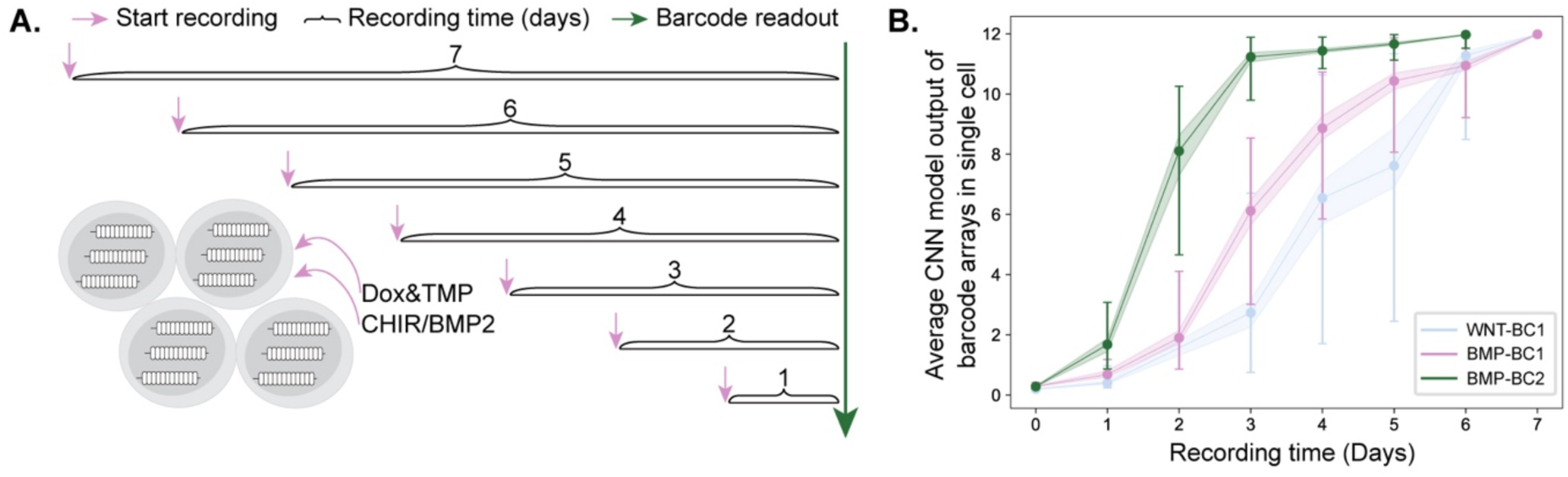
Recording duration of the WNT and BMP signaling in INSCRIBE cells. **(A)** Schematic of experimental workflow for time dependent recording of signaling activity. ABE expression was induced by 500 ng/ml Dox and 10 µM of TMP. The WNT and BMP pathways were activated by addition of 3 µM CHIR and 64 ng/ml BMP2, respectively. Cells were cultured in recording condition for varying amounts of time and processed for ratiometric barcode readout together. **(B)** Population median of single cell edit levels plotted against recording time for each cell line, indicated by color coding. Single cell edit levels were determined by averaging the edit levels of all arrays within the cell, as inferred by the CNN classifier. Bars represent the interquartile ranges and shaded areas show the 95% confidence interval for the medians.

### INSCRIBE barcode arrays are stable and efficiently utilized

Since our barcode arrays contain 12 tandem repeats of identical gRNA target sites, it is important to verify their long term stability in the genome. Collapse of barcode sequences, leading to loss of memory, has been observed in other molecular recording systems ^31^. INSCRIBE uses base editing, which avoids double-strand DNA breaks ^43^, and is therefore expected to be less prone to collapse during editing. To assess the stability and editing patterns of barcodes in INSCRIBE cells, we used long-read amplicon sequencing. Further, we combined amplicon sequencing with imaging to validate ratiometric readout against an independent method **(Fig. 4A)**.

**Figure 4.**
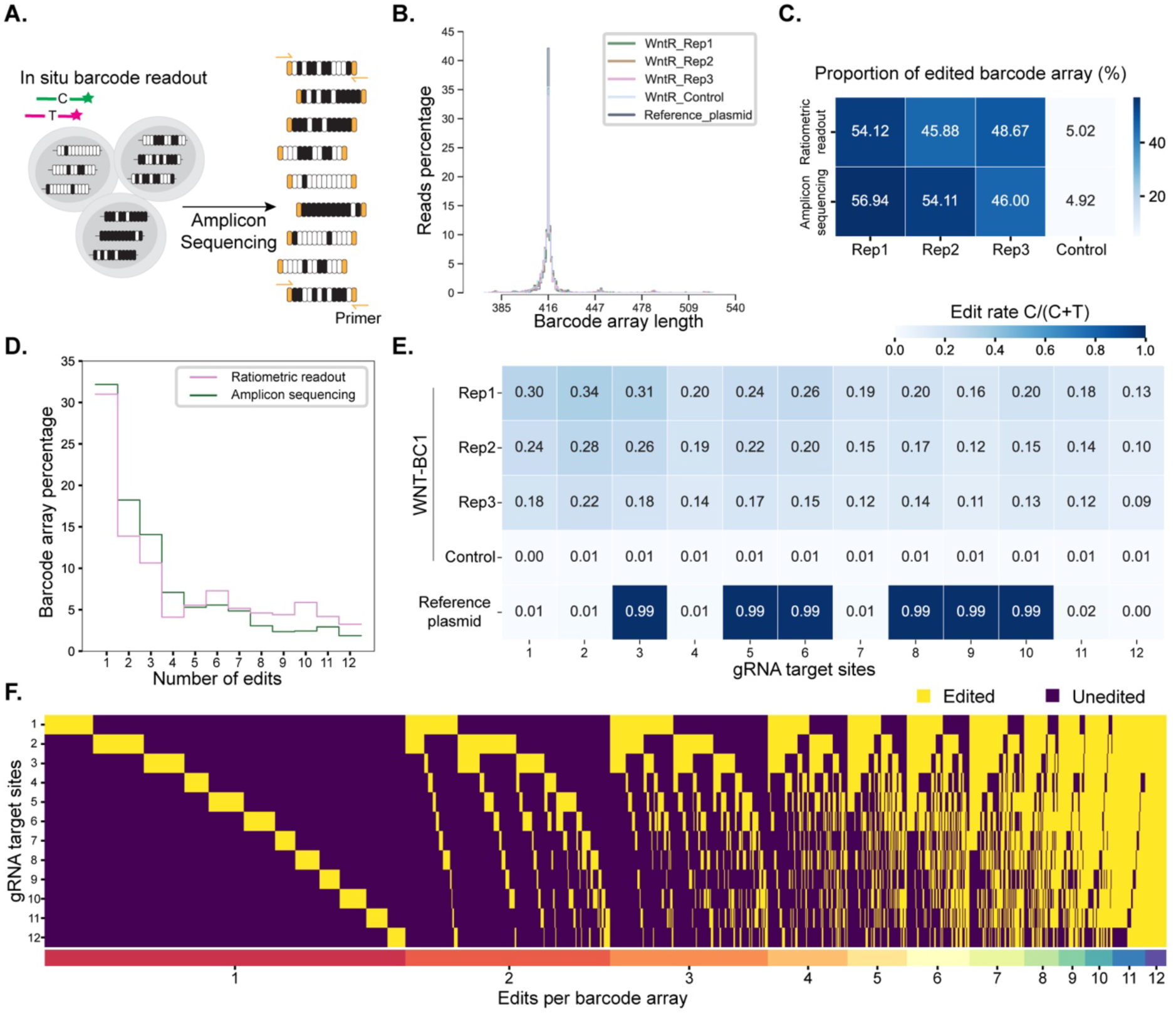
Evaluating genomic stability and edit patterns of INSCRIBE barcode arrays by sequencing. **(A)** WNT-BC1 cells from the same population were subject to imaging-based ratiometric readout and next generation amplicon sequencing. Recording was induced by 500 ng/ml Dox and 10 µM of TMP for 3 days. The WNT pathway was activated during recording with 3 µM CHIR. **(B)** Histogram of barcode array length, recovered from amplicon sequencing, for WNT-BC1 cells after recording (three replicates) as well as an unstimulated control population maintained in culture for over a year. The length distribution for the barcode array amplified from a plasmid is included as a reference. **(C)** Heatmap showing the proportion of edited barcode arrays recovered from ratiometric barcode readout and amplicon sequencing. Edited arrays in imaging-based analysis are identified as those whose edit level based on the CNN model output is above 95th percentile of arrays in unstimulated control condition. **(D)** Histogram of number of edits in edited barcode arrays based on ratiometric readout (pink) and amplicon sequencing (green). For imaging-based analysis, the CNN model output is binned into 12 classes. **(E)** Heatmap of edit level for each gRNA target site in the barcode array recovered from amplicon sequencing. Results of the same analysis for a reference plasmid with known edit status (001011011100) is used to establish the single base pair accuracy of long read amplicon sequencing. **(F)** The edit status of all edited barcode arrays recovered by amplicon sequencing from three replicates of WNT recording experiments.

We amplified barcode arrays from genomic DNA of WNT-BC1 cells maintained in culture without editing for more than one year as well as WNT-BC1 cells grown in editing conditions for 3 days. Additionally, as a control, we included a barcode array with known edit status (001011011100) that was amplified from a reference plasmid **(Fig. S4A)**. For all samples, amplicon sequencing showed the same distribution of array lengths centered on the correct length of 416 bp, confirming the stability of INSCRIBE barcode arrays **(Fig. 4B)**.

Sequencing results were also consistent with *in situ* measurement of barcode edits by ratiometric readout. We first compared the fraction of barcode arrays with at least one edit between sequencing reads and ratiometric readout of cells from the same population. Both methods measured the proportion of edited barcode arrays to be close to 50% in WNT-BC1 cells grown in editing condition for 3 days and close to 5% for the control unstimulated cells **(Fig. 4C)**. Further, within the edited barcode arrays of WNT-BC1 cells after 3 days of recording, the number of edits estimated by *in situ* readout or sequencing showed a similar distribution **(Fig. 4D and S4B-C)**. Together with our analysis of synthetic barcode arrays with known ground truth **(Fig. 1)**, this provides further assurance that ratiometric readout accurately recovers the fraction of edited sites in the arrays.

Estimating the overall signaling activity is most efficient if all target sites within an array have the same probability of being edited. In contrast, if certain sites are more likely to be edited than others, this bias should be accounted for when integrating the results across all barcodes. Since INSCRIBE barcodes have identical sequences and are targeted by the same gRNA, we expect them to have similar edit rates. To test this, we first confirmed that long read amplicon sequencing is accurate at the single nucleotide level, using an amplicon with known edit state that was amplified from the reference plasmid **(Fig. 4E)**. We then quantified edit levels in each of the 12 target sites in barcode arrays of WNT-BC1 cells, with three replicates for the editing condition **(Fig. 4E)**. Edit levels at each site were significantly higher in the editing condition compared to the control (at least 9 times) but, as expected, there was much less variation between sites. Combined with our earlier results **(Fig. 1E and S1C)**, our analysis shows that neither writing nor reading of edits is sensitive to the position of a barcode within an array. We also found that almost all possible combinations of barcode edits are obtained during recording, and no single combination dominates the outcomes **(Fig. 4F)**, further confirming that the memory capacity of INSCRIBE is used efficiently.

### The memory capacity of INSCRIBE cells is sufficient for accurate reconstruction

INSCRIBE cell lines benefit from multiple genomic integrations of barcode arrays, each providing an independent measurement of base editor activity in a cell. To understand how the memory capacity of INSCRIBE cells influences the reconstruction accuracy, we simulated the process of writing and reading edits based on our experimental results. Base editing is a probabilistic process at the single nucleotide level, with each target site having a probability of being edited in unit time. This probability should be proportional to the signaling activity in the cell. We aim to infer this underlying probability in each cell from the observed edit level of barcode arrays. In simulating the editing process based on our empirical findings **(Fig. 4E)**, we assumed that the editing probability is uniform across all sites within a barcode array, leading to a binomial distribution with 12 trials. We varied the editing probability from 0 to 1, in increments of 0.01, to model different levels of signaling activity, and the same editing probability was applied for different simulated barcode arrays within the same cell. Subsequently, we grouped barcode arrays to simulate cells containing 1 to 10 barcode array integrations. To recreate the readout process, we paired each simulated barcode array with a real instance of the BC1 array with the same number of edits, selected randomly from images of cells with known ground truth **(Fig. 1)**. We then used our CNN model to classify those images and recorded the results as the observed edit level of the simulated barcode arrays. The inferred edit probability in each simulated cell was then calculated by maximum likelihood estimation based on all its barcode arrays.

Our results showed a linear correlation between the inferred and true edit probabilities **(Fig. 5A)**. As the number of barcode arrays per cell increases, the correlation between inferred and true edit probabilities is improved, indicating higher reconstruction accuracy **(Fig. 5B)**. However, there are diminishing returns beyond 5 barcode arrays. Therefore, the number of barcode array integrations in WNT and BMP recording INSCRIBE cell lines **(Fig. 5C)** is within the range required for accurate single cell reconstruction of signaling activity.

**Figure 5.**
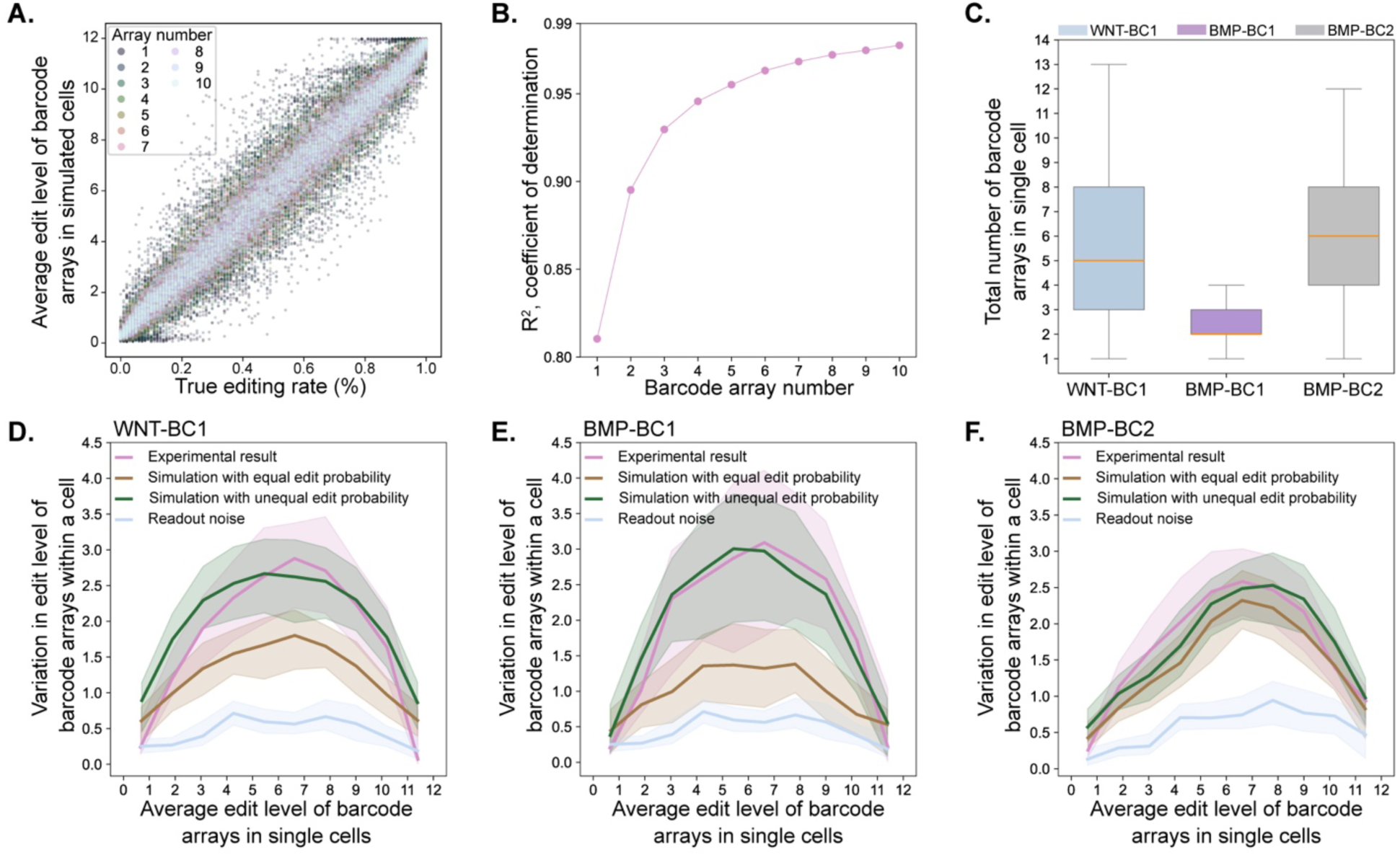
Reconstruction accuracy and noise in signal recording by INSCRIBE. **(A)** Scatter plot shows the single cell inferred edit level against the true edit probability for varying number of barcode arrays, indicated by color coding. 50 cells were simulated for each edit probability and barcode array number. **(B)** The R-squared values for correlation between inferred edit level and true edit probability across different numbers of barcode arrays per cell. **(C)** Distribution of the number of barcode arrays detected in individual cells for each INSCRIBE cell line. Even though each cell line is a monoclonal population, some variability can arise in the number of detected arrays due to efficiency of in situ transcription and imperfect segmentation. Here the data from different recording durations are combined (0 to 7 days for WNT-BC1 and BMP-BC1 and 0 to 6 days for BMP-BC2). WNT-BC1, BMP-BC1, and BMP-BC2 cells were stimulated by 3 µM CHIR, 256 ng/ml BMP2, and 16 ng/ml BMP2, respectively. 500 ng/ml Dox and 10 µM of TMP was used to induce ABE expression. **(D, E and F)** The plots show population median (line) and the interquartile range (shaded area) for the standard deviation against the mean of edit levels of barcode arrays within a cell. Cells with arrays of known edit patterns were used to calculate the readout noise (blue). Experimental results for recording with each INSCRIBE cell line were obtained as described in Fig. 5C (pink). Only cells with more than one array were included in the analysis. Simulations were performed either under the assumption of equal edit probability for all arrays in the same cell (brown) or with added normal noise to vary edit probability for different arrays of a cell (green). The standard deviation of normal noise was 0.21, 0.28, and 0.09 for WNT-BC1 **(D)**, BMP-BC1 **(E)**, and BMP-BC2 **(F)**, respectively.

The integration of multiple barcode arrays can also introduce noise into the recording process, if barcodes at different genomic loci are edited with slightly different rates. To assess this noise, we compared empirical results from each INSCRIBE cell line with simulations similar to those described above **(Fig. 5D-F)**. In each case, we evaluate the variability in recovered edit levels against the mean of barcode edit level in single cells. We first established a baseline for the readout noise, resulting solely from the ratiometric readout process. For this we used measurements of cells with known edit states **(Fig. 1 and Fig. S1)**, where all barcode arrays in a cell have exactly the same number of edits. For any given edit level, variation in recording experiments **(Fig. 5C)** was higher than the baseline readout noise. Part of this additional variation can be attributed to the inherent sampling differences, where each barcode array within a cell may end up with a different number of its 12 barcodes edited, even when all arrays in a cell share the same editing probability. We captured this aspect of the recording noise by simulating cells with integration counts mirroring cells in our experiments and applying the same readout noise. The difference between the variation observed in these simulations and the variation seen in experimental results can be attributed to varying edit probabilities of arrays at different genomic loci. To reflect this, we also performed simulations where a normally distributed noise was added to the cell’s overall edit probability, generating a slightly different edit probability for each array within the cell.

Our analysis showed that, in all cases, the contribution of readout noise to the overall variation is minimal. For WNT-BC1 and BMP-BC1, the empirical variance was consistent with simulations incorporating normally distributed noise with standard deviations of 0.21 and 0.28, respectively **(Fig. 5D-E)**. This suggests that different barcode arrays integrations in these cells are edited at somewhat different rates. In contrast, observed variance in BMP-BC2 cells had considerable overlap with simulations that assume equal edit probability for all barcode arrays **(Fig. 5F)**, and a modest normal noise with standard deviation of 0.09 was sufficient to match the experimental results. Therefore, the positional effect on BC2 appears to be weaker than BC1. This may be due to higher accessibility of BC2 integrations or sequence specific differences between gRNAs.

### INSCRIBE reveals persistent cellular memory in the BMP pathway

Cell-to-cell variability is a common and essential feature of biological systems, allowing for a range of outcomes that influence adaptability and function ^44,45^. Variability is particularly important in the context of signaling, where differences in cellular responses can influence developmental processes and tissue organization. Even in a genetically identical population, individual cells can display a wide range of behaviors and responses to the same external signals ^46^. Origins of this variability can be ascribed to the stochastic nature of protein expression ^47,48^ and regulatory features of the signaling pathways ^49,50^. However, the functional consequences of cellular variability are often difficult to identify, because it requires linking the initial heterogeneity to an eventual phenotype. By recording signaling activity level in the genome of the cells, INSCRIBE provides a way to establish this connection.

Using the INSCRIBE cell lines, we explored whether initial heterogeneity in pathway activity levels results in persistent differences among cells, affecting their likelihood of exhibiting stronger or weaker responses to subsequent stimulations. Specifically, we examined cells exposed to BMP or WNT signals twice, with varying time intervals between exposures **(Fig. 6A)**. Despite being genetically identical, individual cells exhibit variability in their response levels to each stimulation. The initial response was recorded in the genome as barcode edits, which are inherited by progeny cells. Recording was turned off during the second exposure and the magnitude of the second response was measured by H2B-CFP accumulation. We asked if there is a correlation between the magnitudes of response to the first and second stimulations, and if so, how this correlation evolves over time. Correlation in this context signifies a memory of the past response level, which persists over several cell generations.

**Figure 6.**
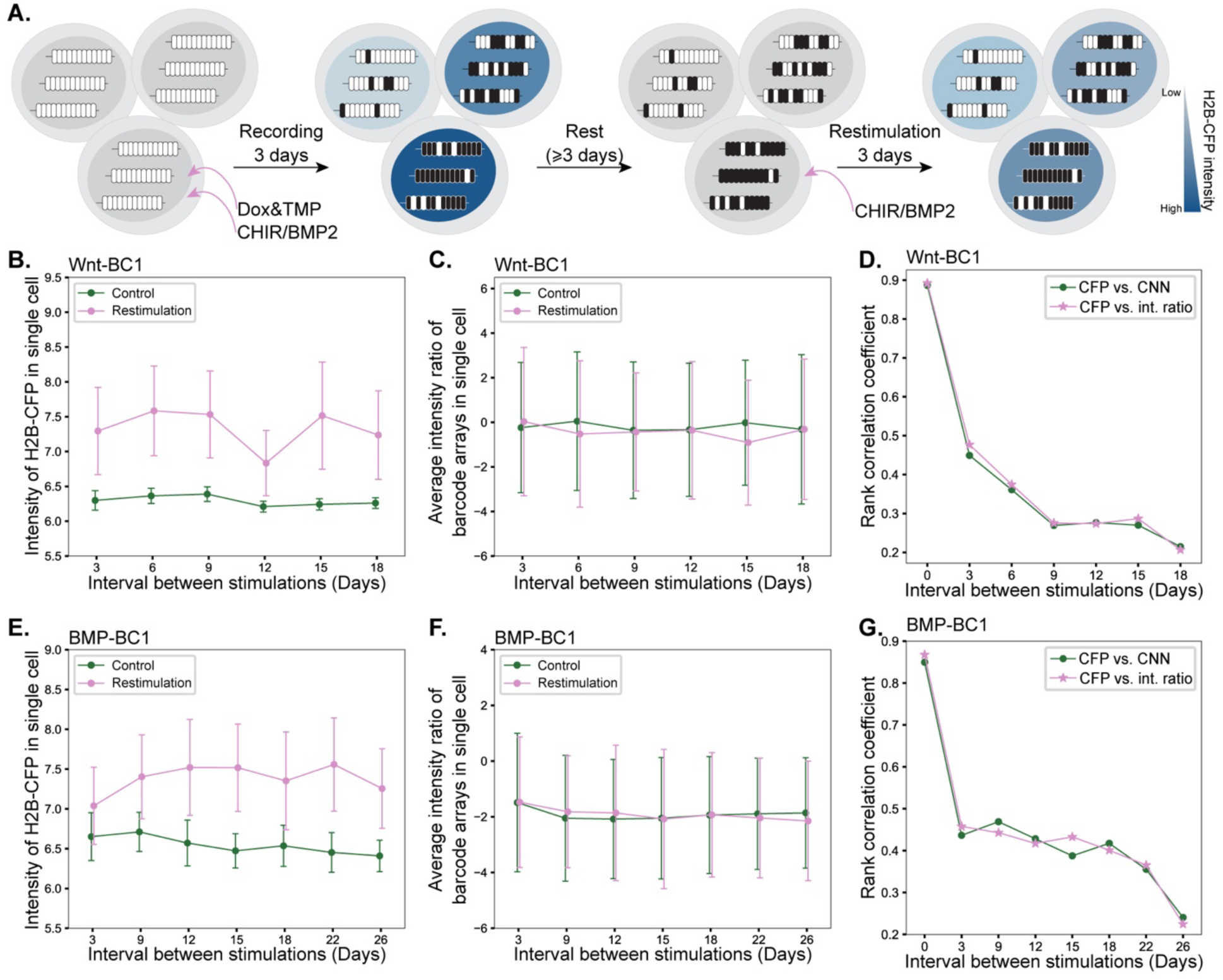
Enduring effects of cell-to-cell variability in response to WNT and BMP signals. **(A)** INSCRIBE cells were exposed to 3 µM CHIR or 256 ng/ml BMP2 to activate the WNT or BMP pathway, respectively, twice with varying time intervals in between. Recording was induced only during the first stimulation, by addition of Dox (100 ng/ml for WNT-BC1 cells and 500 ng/ml for BMP-BC1 cells) and 10 µM TMP. **(B)** Population average (points) and standard deviation (bars) of single cell H2B-CFP intensity after restimulation of the WNT pathway (pink). In the control group (green), cells were stimulated only once during the first 3 days of recording and CHIR was not added to the media during the second stimulation period. The results indicate that H2B-CFP intensity reaches a basal level after the rest period (green). Therefore, the H2B-CFP produced during the first stimulation does not impact the endpoint measurements after restimulation. **(C)** Population average (points) and standard deviation (bars) of single cell edit levels after restimulation of the WNT pathway (pink) and in the control condition (green). The results indicate that recording is off and new edits are not added during the restimulation phase. Edit level was calculated as the average of intensity ratios for all barcode arrays in each cell. **(D)** The rank correlation between the first and second response levels for varying time intervals between stimulations of the WNT pathway. The first response level was measured either by intensity ratio (pink) or the CNN model (green). The second response level was measured by endpoint H2B-CFP intensity. **(E, F, and G)** Similar analysis as in B, C, and D, performed on BMP-BC1 cells through two sequential activations of the BMP pathway by BMP2.

For both pathways, H2B-CFP produced during the first exposure was diluted to a consistent low baseline level within 3 days after the signal was removed **(Fig. 6B and 6E, green data points)**. In contrast, when cells were stimulated a second time, their H2B-CFP levels were elevated above the baseline by the end of the experiment **(Fig. 6B and 6E, pink data points)**, indicating that in our experiments the endpoint CFP intensity reflects only the magnitude of the second response. Similarly, we confirmed that barcode edits capture only the initial response level by demonstrating that edit levels remained unchanged after the second stimulation, as no Dox and TMP were added to induce recording during this phase **(Fig. 6C and 6F)**. While Dox is necessary for the expression of ABE and the accumulation of edits, it is not required for the expression of H2B-CFP from the signal-responsive promoters. This allowed us to measure the amplitude of response to the first and second stimulations separately.

We then examined the correlation between the first and second response levels at the single-cell level. As anticipated from our earlier results **(Fig. 2)**, immediately after treating WNT-BC1 cells with CHIR and BMP-BC1 cells with BMP2 for 3 days, there was a strong correlation between CFP and edit levels inferred from intensity ratio (0.89 and 0.87, respectively) or the CNN model (0.89 and 0.85, respectively). For the WNT pathway, the correlation between the magnitudes of the first and second responses decreased rapidly and consistently as the interval between the two stimulations increased from 3 to 18 days **(Fig. 6D)**. Remarkably, the BMP pathway exhibited different dynamics **(Fig. 6G)**. The correlation remained nearly steady, fluctuating between 0.4 and 0.5, for stimulations up to 18 days apart, and declined only for intervals beyond 3 weeks. This finding reveals that activation of the BMP pathway induces a lasting cellular memory. Descendants of cells that exhibit a strong BMP response tend to maintain an elevated responsiveness to subsequent BMP stimulation, even up to 18 days later. This phenomenon cannot be attributed to genetic differences between cells, as the observed memory effect dissipates by 26 days after the initial exposure.

## Discussion

The fate of a cell is impacted by factors such as lineage and signaling history that are often inaccessible to direct observation. Programming cells to chronicle their molecular history in their genome over time has emerged as a promising solution to this fundamental problem^26–28^. Here we introduce INSCRIBE, a new approach to imaging-based molecular recording that infers past signaling activity in individual cells from endpoint measurements, without the need for sequential imaging. INSCRIBE permanently records the signaling activity level over a specified time, or the duration of signaling with a known intensity, as a set of single base edits in genomic barcode arrays. The fraction of edited sites can be detected at a later time using ratiometric barcode readout, which only requires ordinary fluorescence microscopy with two channels for each signal of interest.

INSCRIBE is made possible through several key innovations. First, the production of gRNA in a signal-dependent manner is enabled by releasing functional gRNA from Pol2 transcripts using HH and HDV ribozymes. Second, an inducible CRISPR base editor introduces precise and predictable mutations at designated sites within the barcode arrays exclusively during the recording phase. Third, ratiometric barcode readout leverages probe competition to infer the number of mutations in each barcode array. Ratiometric readout drastically reduces the cost and technical challenges associated with existing imaging-based barcode readout methods, enabling greater access for researchers. It also cuts down the overall imaging time significantly, and thereby facilitates analysis of more cells and larger samples. Further, *in situ* transcription of barcodes by phage RNA polymerase is compatible with FISH-based spatial genomics methods that enable deep characterization of cell states at endpoint ^22,32^. Fourth, by integrating information across multiple target arrays in each cell, INSCRIBE achieves higher single cell reconstruction accuracy. Finally, we developed a computational pipeline for analyzing barcode images, including CNN models to classify spots corresponding to each barcode array. Along with straightforward and cost-effective experimental procedures, we anticipate this will facilitate the widespread adoption of INSCRIBE for studying how past signaling events shape development and disease outcomes.

We tested INSCRIBE in HEK293 cell lines engineered to record the activity of either BMP or WNT pathway. Barcode edits accumulated in INSCRIBE cells in proportion to the signal level (BMP2 or CHIR concentration) and signaling duration. In contrast with reporter systems that monitor pathway activity by expressing a fluorescent protein, recorder systems like INSCRIBE capture pathway activity as DNA mutations. However, both reporter and recorder assays rely on cis-regulatory elements (CREs) that drive expression of a transcript proportional to the signaling activity. Signaling pathways can elicit complex and multifaceted responses in cells, with various endogenous targets expressed at different levels and exhibiting distinct dynamics^51^. So a single CRE, or a single gene, may not fully capture all the nuances of cellular response to a signal or pathway activity across all cell types. Therefore, the choice of CRE depends on the specifics of the model system and research objective. In principle, INSCRIBE’s recording capabilities are not restricted to transcriptional regulation of gRNA through CREs; any signal that can be converted into a proportional change in edit rate can be recorded and read out similarly. For example, future work could link the edit rate to endogenous protein levels, by fusing Cas9 to conditionally stable nanobodies ^52^.

Using INSCRIBE, we discovered a persistent memory in the BMP pathway, evidenced by the correlation between response levels to two stimulations occurring up to 18 days apart. To our knowledge, this phenomenon has not been previously reported, which opens up intriguing questions regarding its underlying mechanism and biological significance. Developmental signaling pathways, including BMP, are utilized repeatedly to regulate various processes such as growth, differentiation, and morphogenesis throughout different stages of development. We speculate that memory of past signaling activity could serve to coordinate these diverse aspects of tissue development. For instance, an initial signal gradient might direct cell fate decisions, while a later uniform signal promotes proliferation. By retaining the memory of the initial response, the system can fine-tune the population sizes of different cell types, ensuring proper tissue development and organization.

The time scale of this memory suggests an epigenetic mechanism, for example chromatin remodeling that correlates with initial response and keeps loci poised for reactivation in subsequent stimulations in the same cell and its progeny. Certain gene regulatory network motifs, specifically positive autoregulation and positive feedback loops, can also provide a memory of input signals ^53^. Positive autoregulation has been demonstrated for BMP2^54^, BMP4 ^55^ and BMP7 ^55,56^. Positive feedback loop is also suggested to be a conserved feature of BMP signaling pathway across vertebrates ^57^. Thus, the regulatory architecture of the BMP pathway may play a role in the memory effect described here. However, these effects are less likely to be cell-autonomous.

While INSCRIBE offers an innovative solution for recording signaling activity, its current implementation also has several limitations that need to be considered. First, the current implementation of INSCRIBE only records cumulative signaling activity, making it difficult to distinguish between an intense, short-lived signal and a lower-intensity, sustained one. Temporal control of recording using inducible ABE can partially address this issue by allowing recordings over different time windows to provide insights into signaling dynamics. Second, the dynamic range of INSCRIBE must be calibrated to align with the signal intensity and recording duration. We found that different barcode arrays exhibit varying edit rates based on their gRNA sequences; high edit rates are suitable for weak signals or short recording periods, while low edit rates are better for strong signals or longer durations. Another approach is to use a suboptimal gRNA, such as one with a mismatch for its target site, to decrease the edit rate of a given barcode array^58^. Finally, ABE expression may interfere with some cellular processes through gRNA-independent off-target editing of RNA transcripts ^59^. Therefore, newer versions of ABE with reduced off-target RNA-editing activity may be better suited for sensitive applications ^60^.

In summary, INSCRIBE provides a versatile, scalable, and quantitative method for genetic recording of biological signals with in situ readout. The detection of barcode edits is greatly simplified by ratiometric readout, offering a faster, cheaper, and more straightforward approach compared to both sequencing– and imaging-based alternatives. Looking ahead, INSCRIBE is poised to adapt readily to a wide variety of developmental systems and disease models. Orthogonal barcode arrays, like the two developed and characterized here, can be used concurrently to simultaneously record multiple signals. Endpoint analysis of cells can be integrated with spatial multi-omics ^61^ to create enhanced cell atlases that combine signaling history, spatial information, gene expression, chromosomal architecture, and chromatin states. By connecting past molecular events to their future outcomes across cells, tissues, and individuals, INSCRIBE will empower development of predictive models of biological processes and pave the way for new strategies to control and manipulate cell fate.

## Methods

### Design of INSCRIBE barcode arrays

INSCRIBE barcode arrays are made of 12 identical barcodes, each 31 bp long, that are assembled together by Golden Gate assembly^62^. Each barcode includes a 20 bp probe site that partially overlaps with a 20 bp gRNA target site. Each gRNA target site was designed to contain only one editable A nucleotide within the activity window of ABE^33^, to ensure that edit outcomes are pure and predictable. Probe sequences were designed with 50% GC content and predicted Tm between 56 and 60°C. We also avoided recognition sites of certain restriction enzymes (BsaI, BsmBI, BpiI, AarI and XbaI) within the memory arrays to facilitate cloning. Identification of individual barcodes within an array in sequencing data is facilitated by unique 4bp cloning scars flanking each barcode.

### Plasmid construction

The sequence of plasmids, probes, and constructs used in this study are listed in Supplementary table 1. All plasmids were sequence verified by full plasmid sequencing (Plasmidsaurus).

The Dox-inducible system was designed as an all-in-one plasmid with the PiggyBac ITRs and was synthesized by VectorBulider. The ABEMax^34^ (Addgene, 112095) is driven by the tet-responsive promoter (TRE3G). The modified rtTA (Tet-On 3G) is constitutively expressed by EF1A promoter and is followed by self-cleaving P2A peptide and puromycin resistance gene. The degron ^36^ was synthesized as a gBlock by IDT and inserted downstream of ABEMax within the same open reading frame, following a 3 x GGGS linker.

gRNAs targeting all the 12 barcodes of each barcode array were integrated downstream of H2B-CFP coding region, spaced by the triple-helix sequence to stabilize the H2B-CFP mRNA that lacks poly-(A) tail after self-cleavage^63^. The gRNA sequence was flanked by Hammerhead (HH) and Hepatitis Delta Virus (HDV) ribozymes ^41^, to release gRNA from transcripts that also encode H2B-CFP. The triple-helix-gRNA-ribozyme sequences were synthesized as gBlocks by IDT and integrated into the target site through HiFi DNA Assembly Cloning Kit (NEB, M5520). WNT– and BMP-responsive promoters were used to drive the expression of the H2B-CFP-gRNA constructs. These plasmids included hygromycin resistance for subsequent selection.

All 12 barcodes and T3 promoter were synthesized as single strand DNA by IDT, then annealed to make double stranded DNA, and assembled together in one reaction by the Bsal restriction enzymes based Golden Gate Assembly (NEB, E1601). The inserts possess BsaI restriction sites at both ends in the proper orientation, each followed by a 4 base pair sequence to achieve ordered assembly. The handle for inserted barcode sites from 1 to 12 and T3 promoter are TGCC-GCAA, GCAA-ACTA, ACTA-TTAC, TTAC-CAGA, CAGA-TGTG, TGTG-GAGC, GAGC-AGGA, AGGA-ATTC, ATTC-CGAA, CGAA-ATAG, ATAG-AAGG, AAGG-AACT, and AACT-ACCG.

### Cell culture

HEK293 (ATCC, CRL-1573) cells were cultured in DMEM (Gibco, 11960069) supplemented with 10% fetal bovine serum (PEAK, PS-FB4), and 1 x penicillin-streptomycin-L-glutamine (Gibco, 10378016) on polystyrene plates at 37 °C and 5% CO2.

For routine passaging, the growth media was removed, after one brief wash with 1 × PBS (Gibco, 14190250), followed by the addition of 0.05% trypsin (Gibco, 25-300-120) and incubation at 37°C in a 5% CO2 incubator for 5 minutes. Trypsin was then neutralized with growth media at room temperature. The cells were then centrifuged for 5 minutes at 300g, the supernatant was discarded, and the cell pellet was resuspended in fresh growth media. The cells were then reseeded into new tissue culture plates for subsequent cell culture.

### Cell line engineering

INSCRIBE components were integrated over multiple rounds of transfection and selection. For all transfection rounds, HEK293 cells were plated in 24 well plates at a density of 200,000 cells per well one day before transfection, then co-transfected with 400 ng of the constructs to be integrated and 100 ng of the piggyBac transposase together with 2 μl of Lipofectamine 2000 (Invitrogen, 11668027) according to the manufacturer’s instruction. One day after, the transfected cells were replated to new 24 well plates and selection was started with corresponding antibiotics. The first layer of insertion was a series of synthetic barcode arrays, 35 cell lines for barcode array 1, and 34 cell lines for barcode array 2 to mimic the different edit status that may arise during recording. These cell lines were engineered in parallel and selected with 5 µg/mL blasticidin (Gibco, R21001). After the ratiometric barcode readout revealed that the majority of the cells had barcode arrays inserted, we integrated the signal responsive gRNA reporter through a second round of transfection into the polyclonal cell lines with unedited barcode arrays, then selected with 100 µg/mL hygromycin (Thermo Fisher, 10687-010). After verifying the reporter response through ligand stimulation at cell population level, we integrated the TetOn inducible ABEMax through a third round of transfection on the top of the previous two insertions, then selected with 2 µg/mL puromycin (InvivoGen, ant-pr-1). In each round of selection, the engineered cells from 1 well of 24 well plate were expanded to 3 wells of 6 well plate, which usually takes 2 to 3 weeks, then frozen for subsequent experiments. Finally, we used 100 ng/ml of Dox (Sigma, D9891) and 10 µM TMP (Sigma, 92131), together with 3 µM of CHIR (TOCRIS, 4423) or 64 ng/ml of BMP2 (R&D, 335-BM) to stimulate the polyclonal population for 3 days to detect the proportion of cells harboring all INSCRIBE components with ratiometric barcode readout, based on which we determined the total number of monoclonal lines to be screened.

### Establishing monoclonal cell lines by cell sorting

The final verified polyclonal cells were trypsinized by 0.05% trypsin (Gibco, 25-300-120) at 37°C in a 5% CO2 incubator for 5 minutes. Subsequently, trypsin was neutralized with growth media at room temperature. The cells were centrifuged for 5 minutes at 300g, the supernatant was discarded, and the cell pellet was resuspended in 1% BSA (Cell Signaling, 9998S) in PBS. The cell suspension was then filtered through a filter cap into flow cytometry tubes, and stained with 5 μl of 7-Amino-Actinomycin D (7-AAD, BD Pharmingen, 559925) per million cells at 4°C for at least 10 mins. Cell sorting was then performed using a fluorescence-activated cell sorter (BD FACSAria) through the 100 µm nozzle, the 7-AAD negative (the viability) and H2B-CFP negative (no leaky reporter expression) cells were gated and collected as single cell per well into 96-well plates, with 4 plates for each INSCRIBE cell line. Single cells were cultured under 200 µl of standard culture medium at 37°C in a 5% CO2 incubator, and the medium was replaced independently for each well every week. Once the monoclonal lines were recovered and expanded, we used 100 ng/ml of Dox (Sigma, D9891) and 10 µM TMP (Sigma, 92131), together with 3 µM of CHIR (TOCRIS, 4423) or 64 ng/ml of BMP2 (R&D, 335-BM) to stimulate the cells for 3 days. Ratiometric barcode readout was then used to screen the lines.

### Signal recording upon pathway stimulation

Dox (Sigma, D9891) was reconstituted in PBS (Gibco, 14190250) to a final concentration of 1 mg/ml. TMP (Sigma, 92131) was reconstituted in DMSO (Sigma, 276855) to a final concentration of 10 mM. CHIR (TOCRIS, 4423) was reconstituted in DMSO to a final concentration of 10 mM. BMP2 (R&D, 335-BM) was reconstituted in a buffer with 4mM HCl (Fluka, 320331), 0.25% BSA (Cell Signaling, 9998S) to a final concentration of 150 μg/ml. All of these reagents were aliquoted and stored at −20°C, except for BMP2 which was stored at −80°C, thawed immediately before use and diluted with the appropriate culture medium.

For the dosage dependent signal recording, 4000 of INSCRIBE cells were seeded on 96 well plate one day before stimulation, then the medium with various concentrations of ligand and Dox combinations was added to each well, followed by recording for three days. For recording signaling duration, 4000 INSCRIBE cells were seeded on 96 well plate at different time points, followed the recording with constant ligand concentration the next day, followed by ratiometric barcode readout at the same time for different durations of recording to eliminate the batch effect. For the signaling memory experiments, 8000 of INSCRIBE cells were seeded on 96 well plate one day before stimulation, after three days of initial stimulation and recording (with Dox and TMP), cells were split into 2 wells, 8000 cells each, and cultured in media without ligands. After every three days, cells were split into 4 wells, 8000 cells for each. Two wells were used for the three days of the second stimulation (without Dox and TMP) or control. The remaining two wells were maintained in culture without ligands, for the next time point. The ratiometric barcode readout for different time intervals between two stimulations was performed immediately after the second stimulation.

### Ratiometric barcode readout

For ratiometric barcode readout, we seeded the cells on glass-bottom 96-well plates (Cellvis, P96-1.5H-N) that were coated with 20 μg/ml laminin (Biolamina, LN511) at 37°C overnight. The preferred cell density was around 10,000 to 20,000 cells per well, to allow straightforward segmentation while maximizing the number of cells analyzed in each field of view (FOV).

Cells were washed with 1 x PBS (Gibco, 14190250), followed by fixation with a fresh mixture of methanol (Sigma, 494437) and acetic acid (Sigma, A6283) with a 3:1 volume ratio at room temperature for 20 min. After two washes in nuclease-free water (IDT, 11-04-02-01) for 5 min each, the cells were incubated with 50 μl of T3 transcription mix at 37 °C for 3 h, which consisted of 0.5 mM NTP (NEB, N0466S), 0.8 U/μl RNase inhibitor (NEB, M0314S), 5 units/μl T3 RNA polymerase (NEB, M0378S) and 1 x RNAPol reaction buffer (NEB, B9012S). After transcription, cells were fixed with fresh 4% formaldehyde solution (Thermo Scientific, 28906) in PBS for 30 min at room temperature followed by two washes with 2 x SSC (Invitrogen, 15557044) for 5 min each, to remove traces of formaldehyde.

Subsequently, cells were incubated in 50μl of hybridization buffer which consisted of 30% formamide (Invitrogen, AM9344), 10% dextran sulfate (Sigma, D8906), and 2 x SSC at 37 °C for at least 30 min. The cells were then incubated with 4 nM of Alexa Fluor 546 conjugated edited probe (IDT) and 4 nM of Alexa Fluor 647 conjugated unedited probe (IDT) in 50 μl of hybridization buffer for 20 h at 37 °C. Then, the hybridized cells were washed four times (15 min each) at 37 °C with prewarmed wash buffer which consisted of 30% formamide (Invitrogen, AM9344), 0.1% Triton-X 100 (Sigma, T8787) and 2 x SSC to remove excess probes, followed by a brief wash with 4 x SSC. Nuclei were stained with 1 μg/ml DAPI (Thermo Scientific, 62248) in 4 x SSC for 10 min, followed by a wash with 4 x SSC for 5 min. The plate can then be stored in 4 x SSC with 0.02 unit/µl SUPERase RNase inhibitor (Invitrogen, AM2694) at 4°C or imaged immediately. Unless otherwise stated, all reagents were added at 100 μl per well.

### Fluorescence microscopy

Immediately before the imaging, 4 x SSC was replaced with fresh anti-bleaching buffer, which consisted of 10% glucose (Sigma, G7528), 10 mM Trolox (Sigma, 238813), 1:100 diluted catalase (Sigma, C3155), 1 mg/mL glucose oxidase (Sigma, G2133) and 50 mM Tris-HCl pH 8.0 (Invitrogen, 15568025) in 4 x SSC. The cells were imaged on the Zeiss AXIO Observer Z1 inverted fluorescence microscope equipped with a Plan-Apochromat 63x/1.4 oil immersion objective (Zeiss), ORCA-Flash 4.0 V3 digital CMOS camera (Hamamatsu, C13440) and fluorescence lamp illuminators (X-Cite, 120PC Q). Optical sections were captured with a 21-plane z-stack at 0.5 μm intervals to cover 10 μm thickness to ensure recovery of all the barcode arrays at different z-planes for each position. Imaging settings, including the exposure times (10 ms for the DAPI channel, 50 ms for the CFP channel, 300 ms for the 546 channel, and 1000 ms for the 647 channel), were kept the same for all the experiments to facilitate comparison between images. Positions were chosen solely based on the DAPI channel to avoid bias. Immersion oil with a refractive index of 1.518 (Zeiss) was used to minimize spherical aberrations.

### Image processing

#### Images were processed using custom Python scripts

##### Maximum intensity projection

To combine information across optical sections while avoiding out-of-focus slices, we assessed the focus of each z-slice by calculating the variance of the Laplacian of the corresponding DAPI image. We then scaled these values from 0 to 100 and identified the largest stretch of slices where this normalized measure of focus is above 20. Maximum intensity projection was then applied to this stretch of z-slices for all the channels.

##### Segmentation

For cell segmentation, we used the GPU-based ‘cyto’ model of CellPose ^64^, a generalist, deep learning-based cell segmentation algorithm. This was applied only to DAPI-stained nuclei in order to avoid incorrect segmentation of neighboring cells. Cells that intersected the border of the image were excluded from the analysis. For segmentation of barcode array spots, we manually trained classifiers through the interactive pixel classification workflow of Ilastik^65^, based on one randomly picked position out of at least 7 positions for each investigated condition. We then applied the classifier to the whole data set through batch processing. The probability threshold for Ilastik probability masks was set to 0.5. Any barcode array spots identified outside of cell nuclei or with the size below 3 pixels were excluded from the analysis.

##### Intensity measurement

After applying background subtraction with 50 pixels rolling ball radius, the intensity of barcode array spots in each channel was obtained by the integration of pixel intensity values over the segmented area of the spot. To estimate the edit level of each array, we calculate the intensity ratio, defined as the base-2 logarithm of the fluorescence intensity in the edited channel divided by the intensity in the unedited channel. Using this measure, the edit level of each cell was quantified as the average of intensity ratios for all its barcode arrays. The intensity of H2B-CFP was calculated as the natural log of average pixel intensity over the segmented nuclei.

##### CNN classifier training

To train the barcode classifier, we used two-channel fluorescence images of barcode array spots with known edit states. Images containing multiple spots were filtered out. The remaining images were resized, using Python’s skimage package, to match the dimensions of the largest image in the dataset. Intensity values were also normalized by scaling each channel according to the maximum intensity value of that channel across the dataset. The labels were then one-hot encoded into 13 classes corresponding to the number of possible edits. We splitted the dataset into training, validation, and test sets, with 20% of the data reserved for testing. The remaining 80% was split further, allocating 25% for validation (resulting in an effective 60/20/20 split for training, validation, and testing). For data augmentation, we applied rotation, horizontal and vertical flipping, and nearest-neighbor filling on the training images to enhance the model’s robustness. No augmentation was applied to the validation and test data.

The CNN model comprised four convolutional layers, each with L2 regularization with a factor of 0.001 and leaky ReLU activation with an alpha value of 0.001. Except for the first layer, every convolutional layer is followed by 2D Max pooling with a 2×2 window and a dropout layer with a dropout rate of 0.2. The model then flattens the output from convolutional layers to feed into a dense layer, which also utilizes L2 regularization and leaky ReLU activation, followed by a dropout layer with a dropout rate of 0.35. Finally, the output layer consists of a single unit producing 13 output values, corresponding to the probabilities of the barcode array containing between 0 and 12 edits.

To evaluate the accuracy of ratiometric readout **(Fig. 1F and S1D)**, we assigned the edit number with the highest probability to each spot. In contrast, for the recording experiments, we calculated the expected value by averaging the products of each state’s probability and its corresponding edit number (0 to 12).

### Amplicon sequencing

For amplicon sequencing combined with ratiometric barcode array readout procedure, 100,000 of WNT-BC1 INSCRIBE cells were seeded on 24 well plate one day before stimulation, then 500 µl medium with 3 µM CHIR, 500 ng/ml Dox and 10 µM of TMP was added to each well, followed by recording for three days. The cells were then split to plate 12,000 INSCRIBE cells on glass-bottom 96 well plate (Cellvis, P96-1.5H-N) coated with 20 μg/ml laminin (Biolamina, LN511) for ratiometric barcode array readout. The rest of the cells were plated on a new 24 well plate for the amplicon sequencing. The next day, we performed the ratiometric barcode readout and amplicon sequencing for the cells from the same population in parallel.

The genomic DNA was extracted from the WNT-BC1 INSCRIBE cells using the Genomic DNA Purification Kit (NEB, T3010) according to the manufacturer’s instructions. The targeted region was amplified from collected genomic DNA or reference plasmid using high fidelity Herculase II Fusion DNA Polymerase (Agilent, 600675). The input template for PCR was 100 ng of genomic DNA or 1 ng of reference plasmid in a final reaction volume of 50 μl. The samples were incubated for 2 min at 95 °C; 30 s at 95 °C, 30 s at 56 °C and 60 s at 72 °C for 36 cycles; and 10 min at 72 °C. Amplicon libraries contained the barcode arrays with an extra 50 bp on each side and were sequenced by the Plasmidsaurus Premium PCR sequencing service, which is based on nanopore long-read sequencing. Raw FASTQ files were aligned to a FASTA-format reference file containing the expected amplicon sequences. Alignment was performed using the bowtie 2, and subsequent analysis and data visualization were performed with custom Python scripts. We extracted base calls from each aligned read at the base-editor target sites. The unique 4 bp sequences flanking each barcode were used to identify the position of barcodes in the arrays.

## Data availability

All code and processed data required to replicate our analysis is provided through a Github repository (https://github.com/askarylab/INSCRIBE). Raw data are available upon request from the corresponding author.

## Acknowledgements

We thank the members of the Askary laboratory, especially Z. Samadi for helpful discussions and approaches for data analysis. We thank L. Sanchez-Guardado for insightful discussions. We are grateful to M. Elowitz and the Elowitz laboratory, especially M. Tran and D. Chadly, for critical feedback and for sharing plasmids with signal responsive elements. This work was supported by a grant from the National Eye Institute (R00EY031782 to A.A.) and the UCLA Eli and Edythe Broad Center of Regenerative Medicine and Stem Cell Research Transformative Technology Development Pilot Award, including support from The Rose Hills Foundation Innovator Grant Program.

## Supplementary material

**Supplemental Figure 1.**
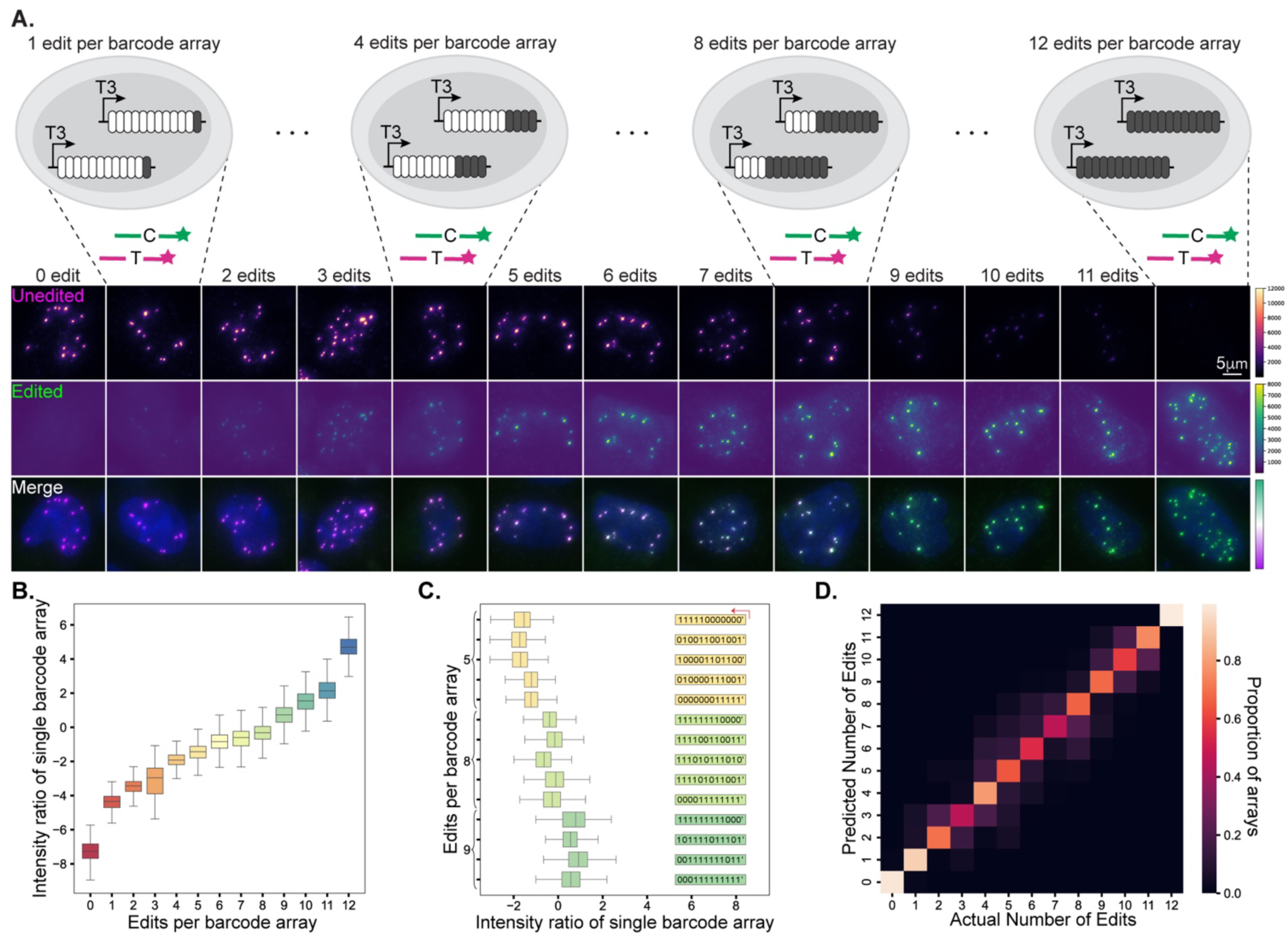
Validation of ratiometric barcode readout for the BC2 array. (**A**) Representative images of cells with the BC2 array with known edit states. Panels are organized in the same way as Fig. 1C. **(B)** Distribution of intensity ratio for each edit number of BC2 array. **(C)** Comparison of arrays with different edit configuration confirms that the number of edits, not their position within the array, determines intensity ratio. **(D)** Classification of BC2 arrays using the CNN model demonstrates accuracy comparable to that achieved for the BC1 arrays. Confusion matrix compares the true edit numbers against the predicted outputs from the CNN model. The frequency values are normalized so that each row sums to one. All results in this figure are from analysis of the BC2 array.

**Supplemental Figure 2.**
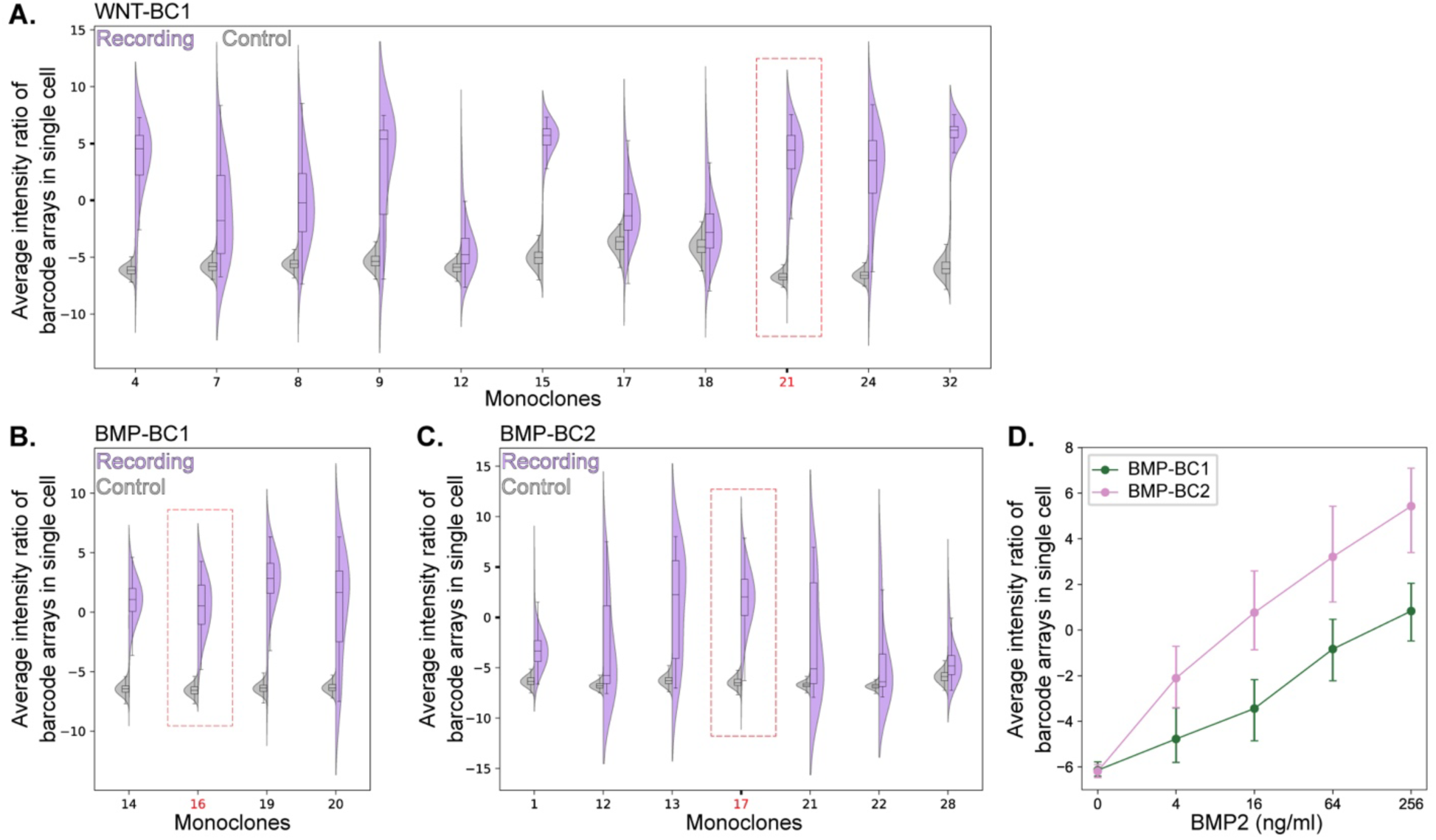
Screening of monoclonal INSCRIBE cell lines. Split violin plots show the distribution of single cell edit levels after pathway stimulation compared to the unstimulated control condition for WNT-BC1 **(A)**, BMP-BC1 **(B)**, and BMP-BC2 **(C)** monoclonal cell lines. Edit level in each cell is calculated as the average of intensity ratios for all its barcode arrays. The WNT pathway was activated by 3 µM CHIR and the BMP pathway was activated by 64 ng/ml BMP2. All the recordings were performed for 3 days, using 100 ng/ml of Dox and 10 µM of TMP. The lines used for further analysis (#21 for WNT-BC1, #16 for BMP-BC1, and #17 for BMP-BC2) are highlighted by red dotted rectangles. **(D)** The population median of single-cell barcode edit levels for BMP-BC1 (green) and BMP-BC2 cells (pink) across different BMP2 levels. Bars represent the interquartile ranges.

**Supplemental Figure 3.**
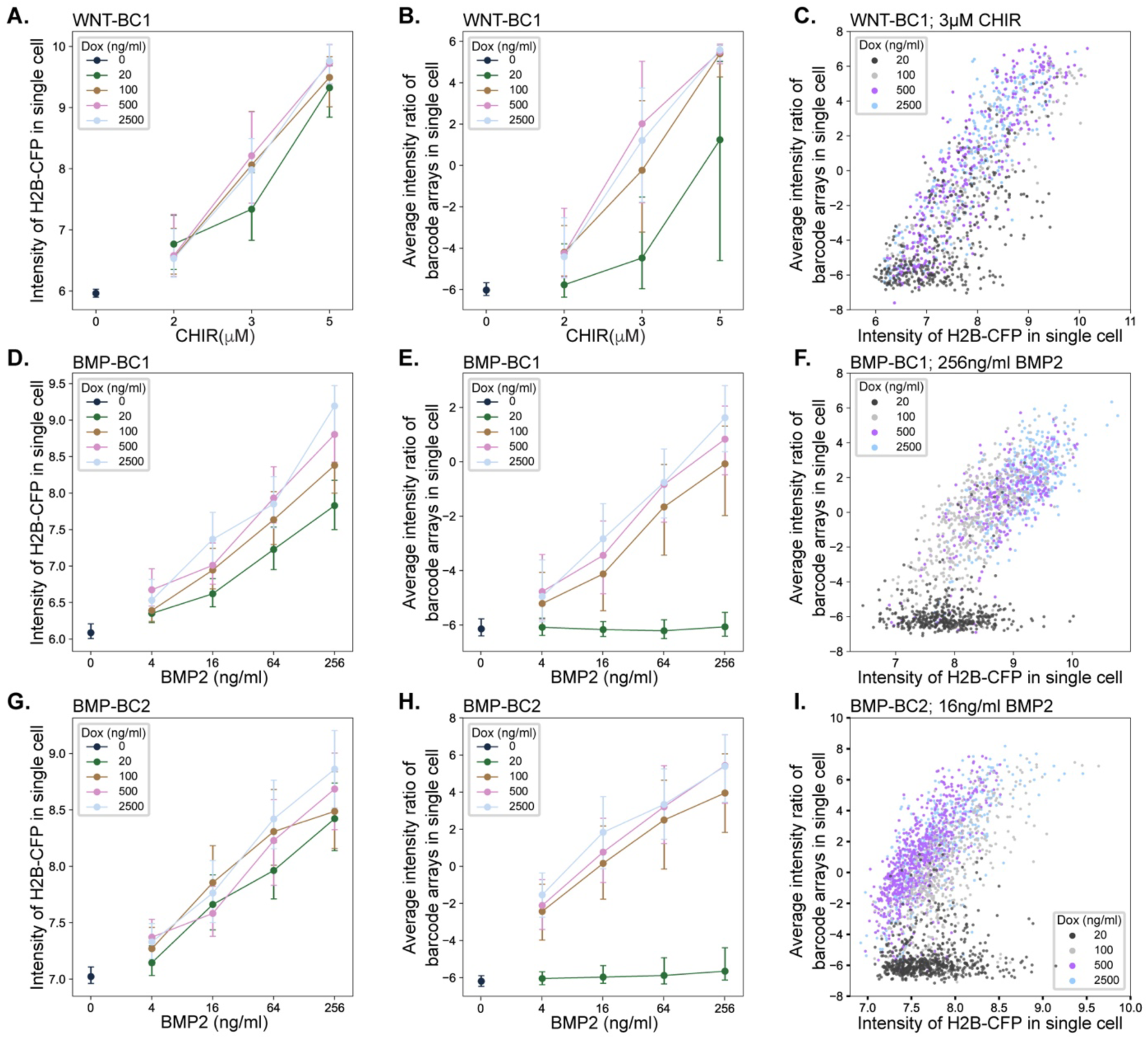
Robustness of signal recording in the INSCRIBE cell lines with respect to the Dox concentration. (**A, B, and C**) Recording of the WNT pathway activity by WNT-BC1 cells. Population medians and the interquartile ranges of single cell H2B-CFP intensities **(A)** and edit levels **(B)** are plotted against CHIR concentration, with colors indicating different Dox levels. While editing requires expression of Dox inducible ABE, H2B-CFP expression is expected to be independent of Dox. The scatter plot **(C)** shows correlation between barcode edit level and H2B-CFP intensity in single cells after 3 days of recording with 3 µM CHIR. Each point in the scatter plots represents a single cell, with colors indicating the Dox concentration. Cells from recording experiments with Dox concentration ranging from 100 to 2500 ng/ml are intermixed. Similar results were obtained for BMP-BC1 cells **(D, E, and F)** and BMP-BC2 cells **(G, H, and I)**. Scatter plots show the results for 3 day recording in the presence of 256 ng/ml BMP2 for BMP-BC1 cells **(F)** and 16 ng/ml BMP2 for BMP-BC2 cells **(I)**. Edit level in each cell is calculated as the average of intensity ratios for all its barcode arrays.

**Supplemental Figure 4.**
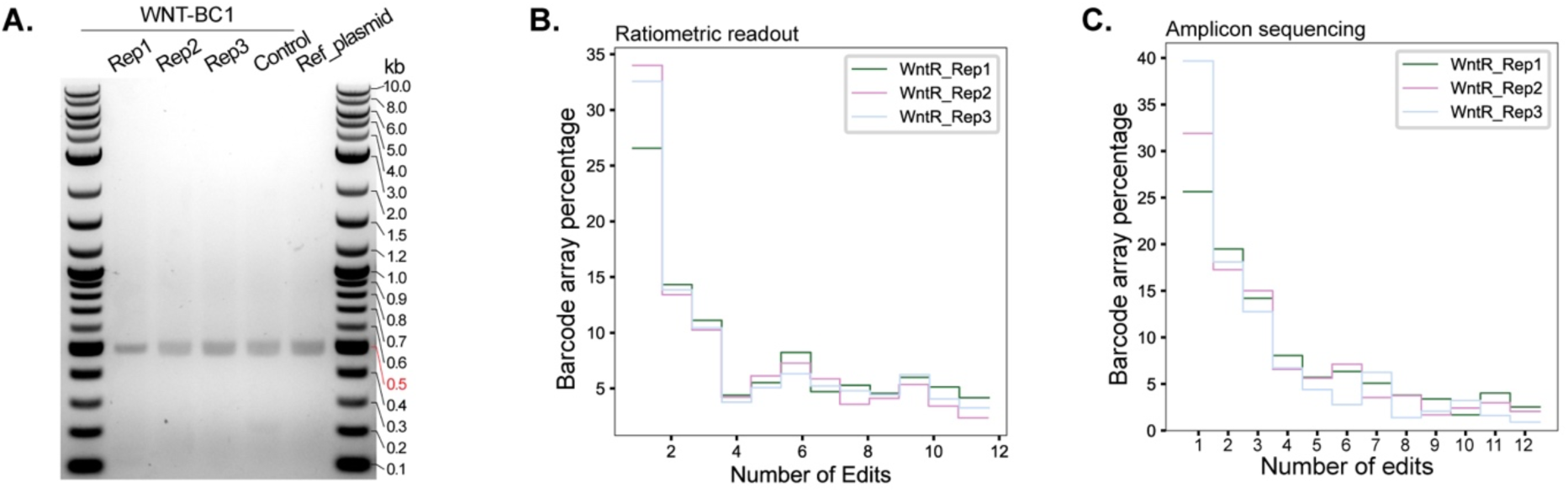
Comparison of number of edits recovered by ratiometric readout and amplicon sequencing. (**A**) Image of DNA electrophoresis gel shows the length of amplicons corresponding with Figure 4B. **(B and C)** Histogram of number of edits in edited barcode arrays based on ratiometric readout **(B)** and amplicon sequencing **(C)** for each replicate of WNT recording experiment.

